# Fabrication of Conductive Hollow Microfibers for Encapsulation of Astrocyte Cells

**DOI:** 10.1101/2022.03.09.483669

**Authors:** Nima Alimoradi, Vahid Nasirian, Saurabh S. Aykar, Marilyn C. McNamara, Amir Ehsan Niaraki-Asli, Reza Montazami, Andrew Makowski, Nicole N. Hashemi

## Abstract

The manufacturing of 3D cell scaffoldings provides advantages for modeling diseases and injuries by physiologically relevant platforms. A triple-flow microfluidic device was developed to rapidly fabricate alginate/graphene hollow microfibers based on the gelation of alginate induced with CaCl_2_. This five-channel pattern actualized continuous mild fabrication of hollow fibers under an optimized flowing rate ratio of 300: 200: 100 μL.min^−1^. The polymer solution was 2.5% alginate in 0.1% graphene, and a 30% polyethylene glycol solution was used as the sheath and core solutions. The morphology and physical properties of microstructures were investigated by scanning electron microscopy, electrochemical, and surface area analyzers. Subsequently, these conductive microfibers’ biocompatibility was studied by encapsulating mouse astrocyte cells within these scaffolds. The cells could successfully survive both the manufacturing process and prolonged encapsulation for up to 8 days. These unique 3D hollow scaffolds could significantly enhance the available surface area for nutrient transport to the cells. In addition, these conductive hollow scaffolds illustrated unique advantages such as 0.728 cm^3^.gr^−1^ porosity and twice more electrical conductivity in comparison to alginate scaffolds. The results confirm the potential of these scaffolds as a microenvironment that supports cell growth.

## 1. Introduction

Tissue engineering as an assimilation of the engineering and biological principles has made significant progress in developing different substitutes such as native tissues for maintenance, repair, regeneration of damaged tissues, and the study of cell-cell interaction ^1, 2^. In this field of research, cellular activities, scaffolds, and growth factors are considered the primary principles involved ^3, 4^. Most microstructure scaffolds employed for tissue assembly benefit abroad range of advantages such as desired geometry, biocompatibility, biodegradability, porosity manner, and desired mechanical properties ^4–16^. These microstructure scaffolds have been fabricated through various techniques such as phase separation, particulate leaching, microfluidics, hydrogels, rapid prototyping, electrospinning, and self-assembly ^17–20^. Alginate 3D hydrogels are one of the most efficient microstructure scaffolds. These alginate 3D hydrogel frameworks are implemented in the most impressively grown areas in biomedical engineering therapy, drug delivery, filtration, channeling, and target delivery of small volumes of liquid to live organisms ^21–25^. This hydrogel is approved by the food and drug administration (FDA) and could be obtained initially from brown seaweed. It can efficiently hold a large volume of water inside its porous cross-linked network as a critical benefit for developing extracellular matrix during cell encapsulation procedure ^26–30^. Among all alginate hydrogels patterns, its fiber designs have attracted more attention with their similar physiological properties to the configuration of fiber proteins ^31^. In addition, hollow alginate fibers could efficiently mimic native tissue properties for continuous nutrient addition and metabolites removal through their considerable surface-to-volume ratio ^32–34^. The continuous fabrication of hollow alginate microfibers ^27, 30, 35–38^ with desired geometries and mechanical properties has been successfully reported using the microfluidic technique as a cost-effective, convenient method. In addition, this pathway could achieve ideal conditions by adjusting the flow rate ratios (FRRs) between the fluid core and sheath flows. The microfluidic hollow microfibers obtained under a gentle polymerization represented a multitude of potentials to be used in various applications such as tissue engineering, cell bioreactors, and biopharmaceutical purification ^27, 35, 39–45^. Recently, the successful induction of the electrical properties to hydrogel structures by conductive biocompatible modifiers such as graphene, graphene oxide, reduced graphene oxide, and synthetic polymers such as polypyrrole or poly(3,4-ethylenedioxythiophene) polystyrene sulfonate (PEDOT:PSS) has been investigated by a number studies to elucidate electrical cell-to-cell communication mechanisms within neuronal cell cultures ^15 46–54^. Graphene owning two-dimensional honey-combed structure of *sp*^2^ hybridized carbons, outstanding biocompatibility, high conductivity, and mechanical properties, has drawn considerable attention in electrophysiology applications compared to other conductive substances ^49, 55–57^. However, such promising nanostructure requires additional mechanical and electronic equipment or some typically toxic surfactants to dominate the interfacial interactions between its carbonic layers for employment in cell-encapsulated hydrogels ^56, 58–60^. A mechanical method based on stirring of graphite in the presence of water-soluble bovine serum albumin (BSA) had been recently reported to produce non-aggregated aqueous graphene solution with high stability ^56, 61–63^. BSA with positive and negative charged spots could successfully lead to the fabrication of aqueous graphene dispersion and be considered an alternative competent for thermal or chemical reduction of graphene oxide that requires extensive use of cytotoxic chemicals to maintain the aqueous graphene dispersions ^56^.

Here, a detailed investigation is provided on the fabrication and characterization of microfluidic hollow fibers obtained from the alginate/BSA-graphene mixture. Furthermore, the potentials of such a biocompatible hydrogel platform for the surviving, regeneration, and electrical stimulation of the human nervous system cells are explained. Our obtained results can pave the way to real-time sensing platforms with control on the cell location and encapsulation.

## 2. Methods

### 2.1. Materials

Dulbecco’s modified Eagle’s medium (DMEM) and Pen/strep solution (penicillin 10,000 U mL^−1^/streptomycin 10,000 μg mL^−1^, 15140-122, 100 mL) were purchased from Gibco Laboratories (Gibco Life Technologies Limited, Paisley, UK) and Gibco Life Technologies, respectively. Fetal bovine serum (FBS) (Qualified One Shot™, Ref#: A31606-01, 50 mL) and Triton X-100 were from Thermo Fisher Scientific (Waltham, MA). Six-well cell culture clusters (Lot# 23314037) were purchased from Costar®. Polyethylene glycol (PEG) (Mn=20000), Graphite (Synthetic graphite powder <20 μm), paraformaldehyde, and BSA (A7906) were purchased from Sigma-Aldrich (St. Louis, MO). The very low viscosity sodium alginate was from Alfa Aesar (Ward Hill, MA, USA). CaCl_2_.2H_2_O was from Fisher Chemical, Waltham, MA, USA. The aqueous solutions were sterilized using a 3 μm pore size polytetrafluoroethylene (PTFE) syringe filter (Tisch Scientific, North Bend, OH, USA), then a 0.45 μm pore size polyvinyl difluoride (PVDF) syringe filter (Fisherbrand, Houston, TX, USA). For all experiments, the ultrapure water (18.2 MΩ cm) was prepared by the Thermo Fisher Scientific water system (Waltham, MA). All other chemicals used were of AR grades and were used without further purification.

### 2.2. Instruments

A JCM-6000 NeoScope Benchtop scanning electron microscope (SEM, JEOL Ltd, Japan) at 15 kV acceleration voltage with a secondary electron detector was used to study the morphology of air-dried structure obtained. A Zeiss Axio Observer Z1 Inverted Microscope (Carl Zeiss, Oberkochen, Germany) was used to capture Fluorescent images. Image processing was carried out with AxioVision Special Edition 64-bit software. The electrical behaviors of the hollow microfibers were determined by Electrochemical Impedance Spectroscopy (EIS), Cyclic Voltammetry (CV), and Galvanostatic Charge/Discharge (GCD) and using a Potentiostat system (Versa STAT 4, Princeton Applied Research, Princeton, USA). Three GenieTouch™ syringe pumps (Kent Scientific Corporation) were used to inject the solutions. A 4-axis CNC USB controller Mk3/4 for mini CNC mill was used to mill the five-channel microfluidic device controlled by a PlanetCNC® (Ljubljana, Slovenia).

### 2.3. Fabrication of Microfluidic Devices

The microfluidic devices used in this study were fabricated from 6.0 mm thickness poly (methyl methacrylate) (PMMA, Grainger, IL, US) using a computerized numerical control (CNC) mini-mill (Minitech Machinery Corporation, Norcross, GA) to mill the core channels and the chevrons with the dimension of 1.00 mm × 0.75 mm (width × height) and 0.375 mm × 0.25 mm (width × height), respectively. For this aim, the two faces of the PMMA chip were milled separately and then bonded together to obtain a microfluidic device. The employed AlTiN-coated end mill cutters and drill bits were purchased from Harvey tools and Grainger.

### 2.4. Preparation of Soluble Graphene Samples through Ball Milling Process

The aqueous BSA-graphene was used to enhance this pre-hydrogel solution’s electrical conductivity. The few-layer graphene (FLG) was fabricated through the liquid-phase exfoliation procedure of the graphite crystallites, approximately 20 μm in size. For this aim, an aqueous mixture containing 20.00 mg mL-1 graphite and 2.00 mg mL-1 BSA was prepared in plastic containers sealed with glue before placing them in metal containers. Steel balls with the diameter of 11/32” and 1/2” were used to apply for shear tensions at 300 rpm rotational speed. The ratio of the overall balls surface area for all solutions was constant at 500 ± 10 m^2^/m^3^ with respect to the solution volume. The exfoliation process was continued for 90 h shaking.

### 2.5. Cell Culturing

A solution containing 45.0 mL DMEM maintenance media, 5.0 mL FBS, and 0.5 mL penicillin (10,000 U mL^−1^)-streptomycin (10,000 μg mL^−1^) was used for astrocyte C8D1A cells culturing in T-25 flasks while maintaining at 37°C, and under 5% CO_2_ atmosphere. After 70% confluency, these cells were passaged three times before the encapsulation procedure. They were trypsinized by 2.00 mL trypsin solution, and then 1000.0 μL of this obtained cell suspension was added to the 3.0 mL alginate/graphene solution. C8D1A cells were cultured for five days in vitro to provide sufficient time to grow and proliferate within a 3D hydrogel alginate/graphene hollow microfiber.

### 2.6. Preparation of Solutions

During our experiments, 2.5 % alginate solution containing 0.1% BSA-graphene was found with ideal viscosity to resist shear force within the microfluidic channel and fabricate smoother alginate/graphene hollow microfibers. Hence, 0.25 gr sodium alginate powder (Alfa Aesar, Ward Hill, MA) was sterilized by 70% ethanol under a UV lamp and dissolved in 8.0 mL of WFI-Quality Cell Culture grade water (Corning, Corning, NY). In the next step, 1.0 mL of freshly mixed UV-sterilized 0.01 g.L^−1^ BSA-graphene solution was added to the solution mentioned above, and stirring was continued. This obtained polymer solution was mixed with 1.0 mL of cell suspension (3.1725 × 10^6^ cells mL^−1^). 30% polyethylene glycol (PEG) (Aldrich Chemistry, St. Louis, MO) was employed for the sheath and core solutions. As the collection bath solution, a sterilized 30.0 % CaCl_2_.2H_2_O solution was used.

### 2.7. Microfluidic Manufacturing of Alginate/graphene hollow Microfiber

At first, the five-channel microfluidic device and other items were carefully wiped with a tissue containing 70% ethanol and irradiated by UV under a laminar flow hood to ensure sterile conditions during our experiments. The 2.5% alginate sodium solution containing 0.1% graphene was mixed by 3.1725 × 10^6^ C8D1A cells mL^−1^within a sterilized Falcon™ 15 mL conical centrifuge tube. The resulting mixture was placed into two 3.0 mL BD syringes to connect the microfluidic device’s two side polymer channels. Subsequently, the core and sheath solutions were connected into the center channel and the other two side channels to guide and solidify the polymer solution. A core/polymer/sheath FRR of 300:200:100 (μL.min^−1^) rate was applied to flow inside our microfluidic channels. This polymeric mixture was introduced into a 30.0% CaCl2.2H2O collection bath upon exiting the microfluidic device. As a result of the cross-linking procedure between Alginate carboxylate groups, the alginate/graphene hollow microfibers could be solidified in the presence of Ca^2+^ which caused further enhancement of their strength ^27, 64^. The polymerized alginate/graphene hollow microfibers were gathered by a tweezer then transferred into 1X PBS solution to be rinsed. The core regions (30% PEG) were dissolved by this aqueous PBS solution, whereas polymerized areas could be retained the permanent structure. The alginate/graphene hollow microfibers containing C8D1A cells were then maintained for eight days in DMEM media at 37 °C in 5% CO_2_/95% humidified air atmosphere for more cell culturing.

### 2.8. Electrical Characterization

A few cell-free alginate/graphene hollow microfibers were mounted on the top of a polystyrene sheet as a nonconductive flat plate. These microfibers were held on the plate surface by carbon and copper tapes which provided an appropriate electronic connection between the alligator electrode clip and the microfibers heads. The electrical resistance of the fibers was measured by a potentiostat/galvanostat instrument, using the cyclic voltammetry method from −0.1V to +0.1V.

### 2.9. Live-Dead Cell Assays

For the live-dead C8D1A cell monitoring, the Alginate/graphene hollow microfibers containing C8D1A cells were carefully rinsed with FBS-free DMEM media three times. In the next step, 4.0 μL cell Tracker Green 5-chloromethylfluorescein diacetate (CMFDA, 10 μM; Thermo Fisher Scientific, Inc.) and 16.0 μL propidium iodide (PI, 8 μM, Invitrogen, Carlsbad, CA) were mixed with 4.0 mL fresh media and added to prepared samples. After incubation of these treated samples for 30 min at 37°C under a 5% CO_2_ atmosphere, the employed dye solution was replaced with 4.0 mL FBS-DMEM media to suspend the hollow fibers and keep their sample humidity during imaging.

### 2.10. Statistical Analysis

Statistical analysis was carried out with R Project Statistical Software to conduct an Analysis of Variance (ANOVA) to compare the means across samples.

## 3. Results

This study investigated the polymerization of alginate solution doped with aqueous BSA-graphene dispersion within the microfluidic device. Several physical properties of the hollow fibers were obtained by investigating the following parameters; the concentration of CaCl_2_.2H_2_O as the water bath, FRR, density of polymer samples, BSA/graphene concentrations, concentration of core and sheath solution effect on the bonds strength and cross-linking density formed between the alginate’s functional groups. Scheme 1 is a schematic of the microfluidic process to procedure alginate/graphene hollow microfibers involving living C8D1A cells.

**Scheme 1.**
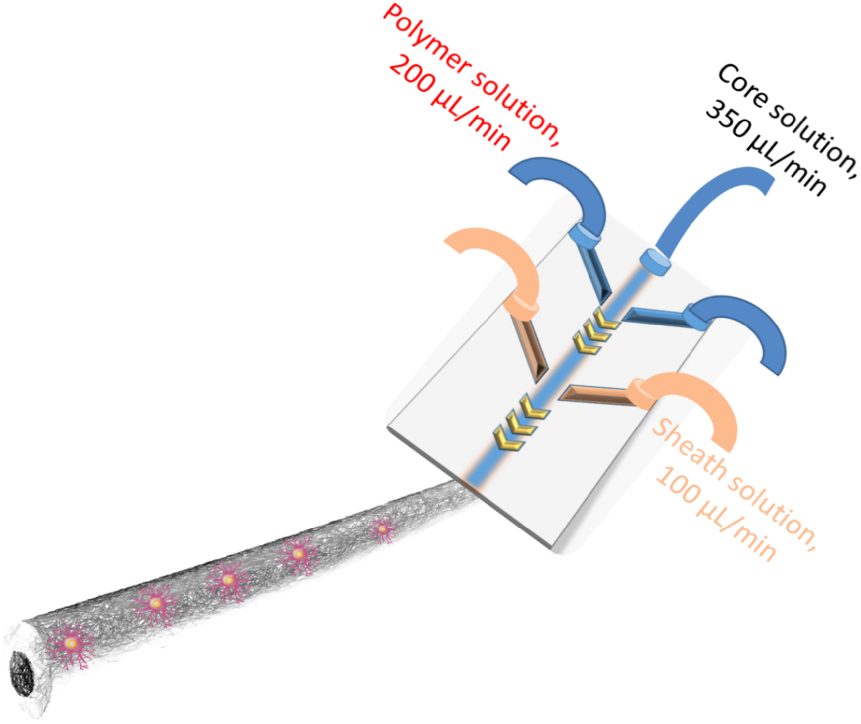
Schematic of Alginate/graphene hollow microfiber generation using a five-channel microfluidic device.

To produce micron-sized hollow microfibers, the polymer, core, and sheath solutions involved in this manufacturing were injected through the channels under the thrusting pump. 30% (w/v) PEG solution was chosen as the core and sheath solutions. This polymer dispersion has high biocompatibility with more cells and so, used extensively in tissue engineering with no significant side effects on cells. Once the PEG has been removed from this architecture by dissolving in CaCl_2_ solution, the hollow channel could be created inside the shell structure to provide the transferring pathway of nutrients and other necessary agents for C8D1A cells into the interior of the structure. The direct relationship between the FRR and the applied hydrodynamic focusing could eventually determine the diameter of obtained hollow fibers. The effect of the influence flow rate ratio on the geometries of these manufactured alginate/graphene hollow microfibers was evaluated, and detailed dimensions of alginate/graphene hollow microfibers under different flow rate ratios are shown in Fig. 1.

**Fig. 1.**
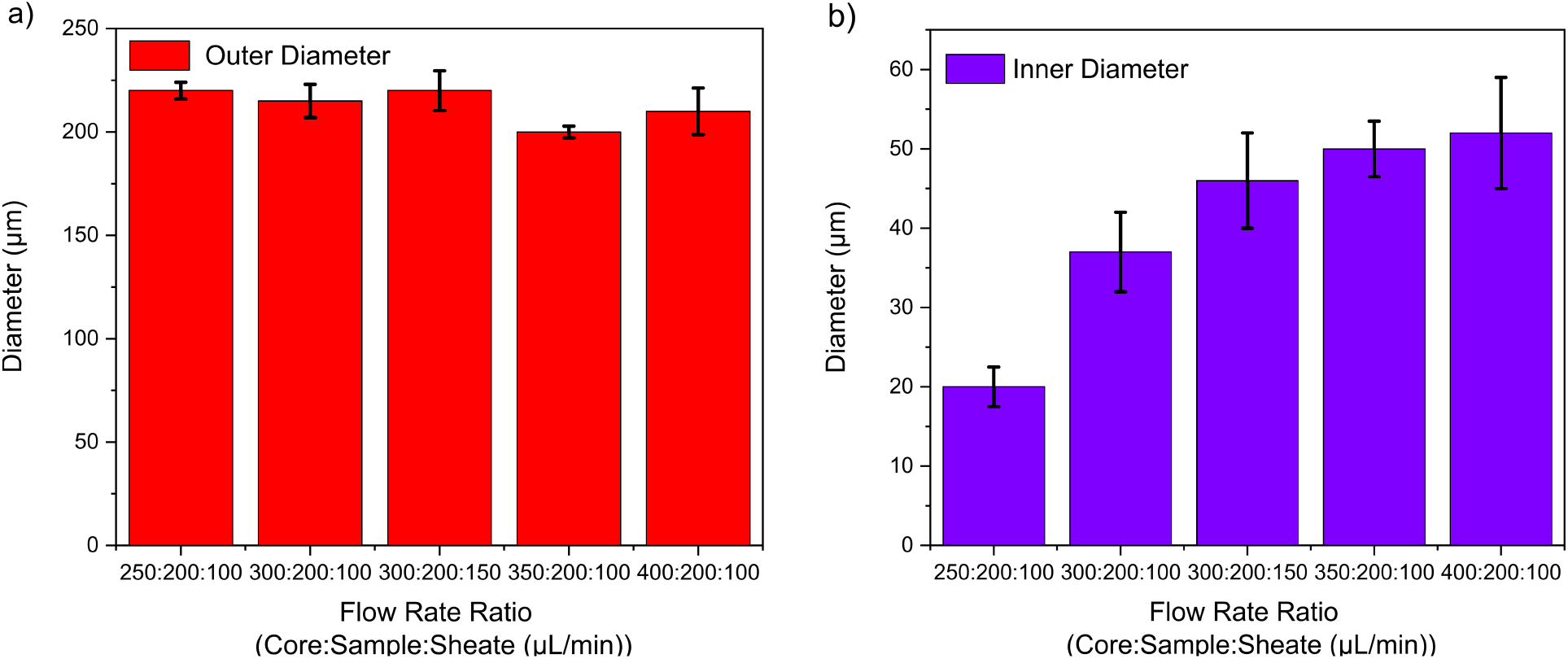
The effect of flow rate ratio on the outer (a) and inner hollow (b) diameters of Alginate/graphene hollow microfibers prepared by the five-channel microfluidic device.

As seen in Fig. 1, the size of these hollow microfibers could be increased more at a lower flow rate ratio due to an enhanced hydrodynamic focusing applied on the core fluid by the polymer and sheath fluids that could lead to loss of cavity diameter ^65^.

### 3.1. Morphology

The microscopic morphology of alginate/graphene hollow microfibers was studied by SEM to determine the sizes and other physical properties of these prepared samples. Fig. 2.a shows the cross-sectional SEM images of manufactured conductive microfluidic hollow microfibers. As seen, the alginate/graphene composite hollow microfibers surface represented a high roughness manner significantly with the average inner-dimension value of about 50.0 μm, which could come from the turbulence that occurred due to the introduction of BSA-graphene to alginate solution. In other words, both the core and edge of alginate/graphene hollow microfiber displayed a porous manner, distributed along the whole structure with sufficient connectivity. By optimizing microfluidic parameters, uniform hollow microfibers that used a 2.5% (w/v) alginate concentration containing 0.1% (w/v) BSA-graphene with an average diameter of 200.0 μm were fabricated. Fig. 2.b shows optical microscope images of the hollow microfibers fabricated when the flow rates for the polymer, sheath and core solutions were maintained at 200, 100, and 350 μl/min, respectively, as optimized parameters.

**Fig. 2.**
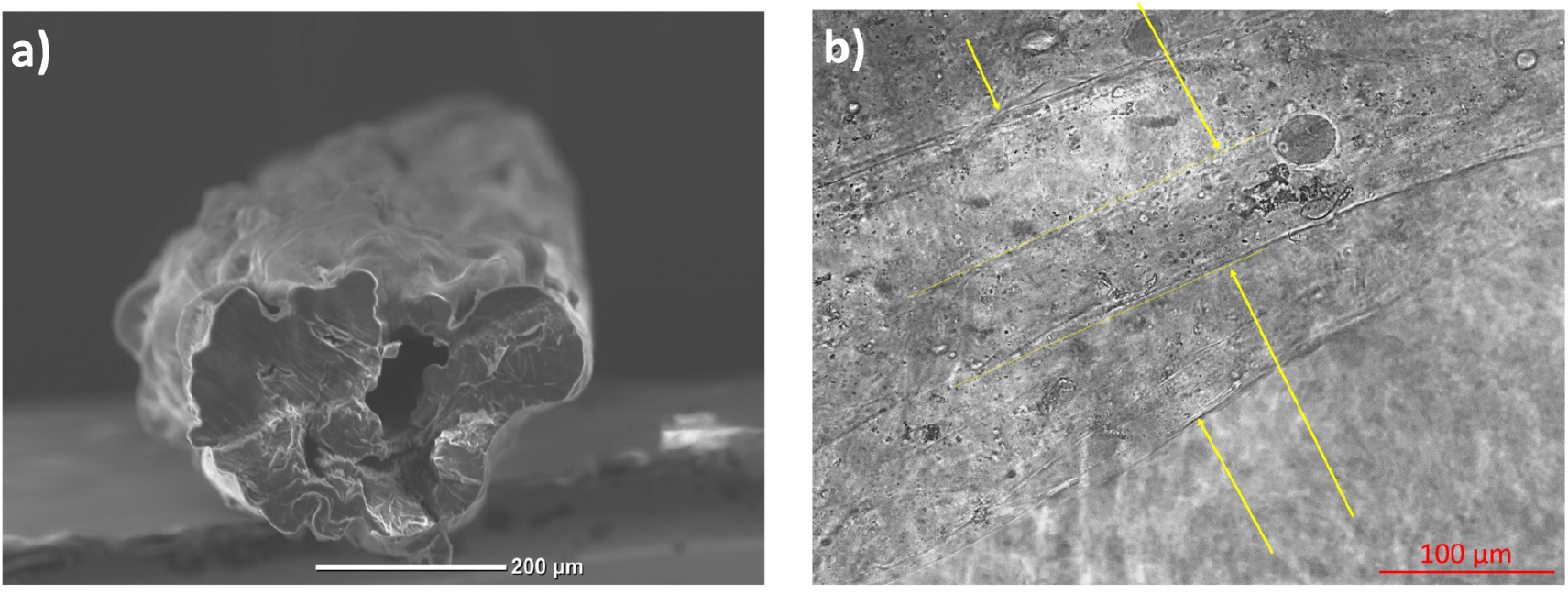
SEM analysis of microfluidic alginate/graphene hollow microfibers manufactured under optimized parameters (a). Illustrative environmental optical microscope images of the alginate/graphene hollow microfibers prepared in this study (b). All microfibers comprised a wall thickness of around 70-90 μm with lumen diameters of ~50 μm.

This (Fig. 2) could successfully confirm the availability of this intermediate duct along the length of these alginate/graphene hollow microfibers with minimal changes in its thickness.

### 3.2. Investigation of Alginate/graphene hollow Microfibers Porosity and Electrochemical Behavior

The electrochemical performance of prepared alginate/graphene hollow microfibers was studied by cyclic voltammetry (CV) and using an electrode constructed by manufactured hollow microfibers and H_3_PO_4_/PVA electrolyte. The CV results obtained were recorded in the range voltage from −0.1 to +0.1 V. the CV curves of prepared alginate/graphene hollow microfibers display a linear behavior which confirmed the effective ion transport throughout the electrode (Fig. 3. a). As expected, a better conductivity of alginate/graphene hollow microfibers with symmetric shape was achieved compared to pure alginate samples. Fig. 3b shows discharge curves regarding the calculated specific capacitance variation of alginate and alginate/graphene hollow microfibers. At the current density range of 0.0-10.0 mA cm^−2^, alginate/graphene hollow microfibers expressed larger specific capacitances than pure hollow alginate microfibers. It is worthy to note that these considerable specific capacitance results obtained for microfluidic alginate/graphene hollow microfibers could successfully confirm the manufactured uniform porous network after introducing graphene to alginate. Moreover, alginate/graphene hollow microfibers can still preserve their substantial capacitance when the current density was increased to 10.0 mA cm^−2^.

**Fig. 3.**
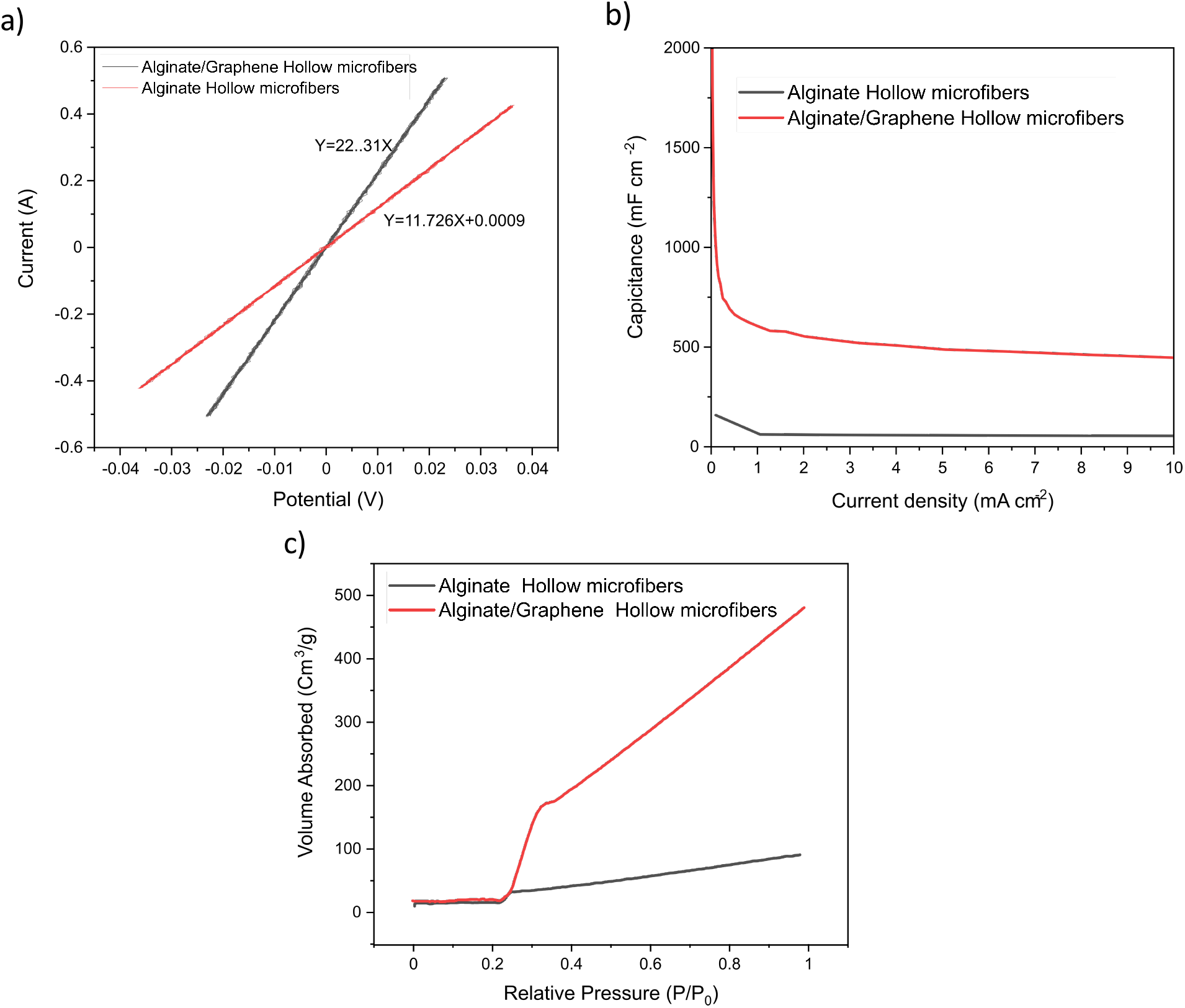
Electrochemical and porosity characterization: CV curves of alginate and alginate/graphene hollow microfiber at a scan rate of 100.0 mV s^−1^ (a). The specific capacitances alginate and alginate/graphene hollow microfiber under different current densities (b). The porosities of microfluidic alginate, and alginate/graphene hollow microfibers, the inset is typical nitrogen adsorption and desorption isotherms of hollow microfibers (c).

The porosity of microfluidic hollow microfibers is another character that must be investigated as the most advantageous parameter when these scaffolds will be used for cell culture inside their lumen. In this case, the cage obtained from the alginate–Ca–alginate linkages could efficiently prepare the crucial room for constituting the mesoporous structure of microfluidic alginate/graphene hollow microfibers walls. On the other hand, these prospered hollow microfibers could express high hydrophilic behaviors, which allowed the media to permeate efficiently through the alginate/graphene wall. These illustrious properties of alginate/graphene and hollow alginate microfibers were successfully confirmed by N_2_ absorption/desorption isotherm measurement (Fig. 3 c).

As seen in Fig. 3 c, the total pore volume (*TPV*) of alginate/graphene hollow microfiber surface was obtained to be 0.728 Cm^3^g^−1^, which has been calculated by Eq. 1 as follows:

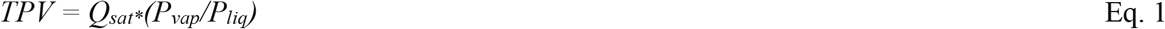

Where Q_sat_ is the N_2_ adsorption quantity, P_vap_ is the density of N_2_ vapor at STP (1.2504 gL^−1^), and P_liq_ is the density of liquid N_2_ at its boiling point (807 gL^−1^). Compared to pure alginate hollow microfiber with TPV of about 0.139 Cm^3^g^−1^, the alginate/graphene hollow microstructure maintained a considerably larger N_2_ volume due to many pores on the alginate/graphene hollow microfiber surface prepared after the addition of graphene.

### 3.3. Immunocytochemistry

As the significant glial cell of the central nervous system, astrocyte cells contain characteristic intermediate glial filament polymers called glial fibrillary acidic protein (GFAP), which are used for astrocyte identification (Fig. 4). For this proceeding, astrocyte cells were seeded on a clean glass coverslip with 30%-40% coverage. When they reached 80-90% coverage, they were washed quickly with 1000 μl of 1XPBS three times. The cells were fixed with fresh-made 4% paraformaldehyde in 1X PBS for 10 min in the next step. Subsequently, wash coverslips with 1X PBS 3 times for 5 minutes each. After, the cells were permeabilized with 0.1% to 0.5% Triton X-100 in 1X PBS for 10-30 minutes. The cells were treated with the blocking buffer (10% FBS, 1% BSA in 1XPBS) for 60 minutes. The cells were incubated overnight at 4°C under a blocking buffer containing the monoclonal Anti-GFAP Alexa Fluor® 488antibody (1:100, Cat #53-9892-82; Thermo Fisher, Waltham, MA, USA). Before taking the images, they were washed with 0.1% Tween 20 in 1XPBS 3 times for 10 minutes.

**Fig. 4.**
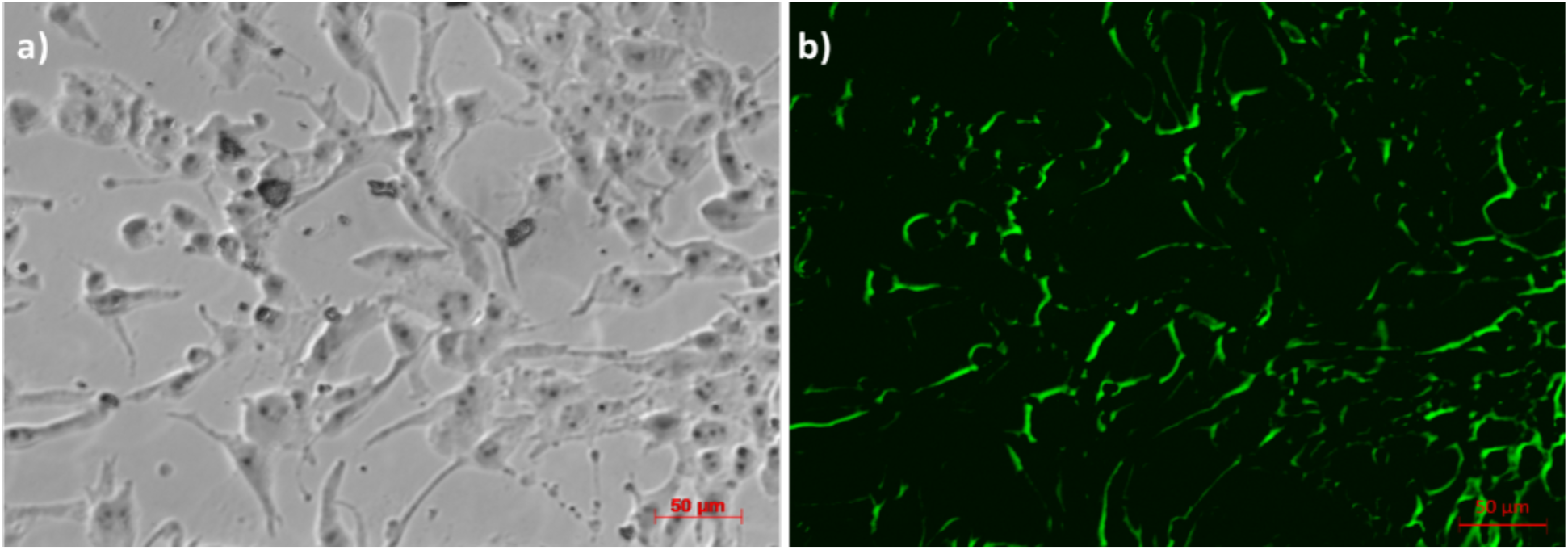
The intermediate filaments of C8D1A were visualized with antibodies against GFAP (green) under excitation wavelength 495 nm, which reveals hypertrophy of cellular processes of reactive astrocytes (b) compared to those under visible light (a).

### 3.4. Cell Viability

Compared to other techniques such as electrospinning as a harsh method that must be operated under high voltage to achieve polymerization, the encapsulation of C8D1A cells within alginate/graphene hollow microfibers using a microfluidic platform can be considered an impressive strategy for the development of biomaterial-based therapeutic. A solution containing 5-Chloromethylfluorescein diacetate (Green-CMFDA) and Propidium Iodide (PI) were employed to study the C8D1A cell viability in this construction and to investigate the 3D environment’s long-term biocompatibility and potential for the influenced C8D1A proliferation and differentiation. Our previous study has shown that the addition of graphene does not change the long-term cell viability; however, gene expression could be altered when cells are in contact with graphene. As a fluorescent nuclear and chromosome counterstain commonly, PI is used to identify the membrane impairment found in dead cells. CMFDA can freely pass through cell membranes and reacts with live cellular components. During these reactions, CMFDA is converted to cell-impartment products, which could pass to daughter cells through several generations without transferring to other cells in the population. Fig. 5 shows the related fluorescence microscopy images taken during 8 days. As seen, the encapsulated C8D1A cells cultured inside the hollow microfibers showed acceptable survival during the microfluidic fabrication process. Furthermore, this cell delivery via hydrogels pathway platform could successfully maintain its confirmation for more cell-cell interactions and the feasibility for an acceptable promising approach in vitro cell culture and regenerative medicine.

**Fig. 5.**
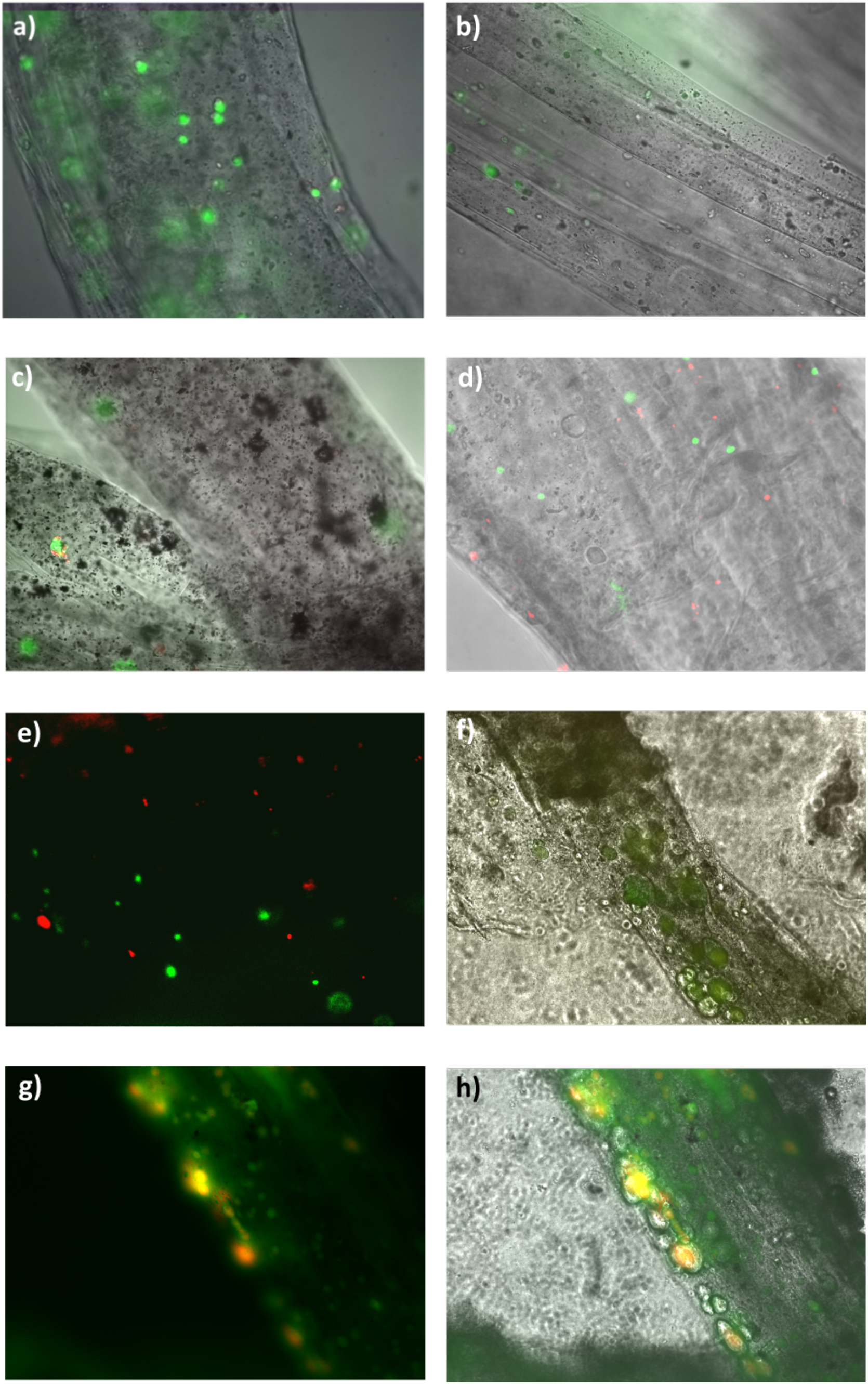
C8D1A cells encapsulated within microfluidic 2.5% Alginate-0.1% Graphene hollow microfibers created with a Core and sheath of 30% PEG and an FRR of 200:300:100 μL min^−1^:μL min^−1^. Samples were gathered in a collection bath of 30% CaCl_2_.2H_2_O. A live-dead assay containing Cell Tracker™ CMFDA (green, live cells) and PI (red, dead cells) was performed on days 1-8 (a-h).

## 4. Conclusions

Altogether, we have developed a tunable alginate/graphene hollow microfiber-based microfluidic platform for long-term supporting in vitro culture of encapsulated C8D1A cells. By introducing BSA-graphene into hollow alginate microfibers, their conductivities are significantly enhanced to deliver and receive electrical signals to cells by a factor of 2. The present findings displayed the first description of porous conductive hollow alginate microfibers to enhance the essential nutrition and metabolic waste transferring between the internal and the external environment of cells. In this study, C8D1A cells were encapsulated in alginate/graphene hollow microfibers as a proof of concept to show that these cells could survive and proliferate in the fabricated microenvironment. We propose further developing this microfluidic technique to achieve a controlled cell number on alginate/graphene hollow microfibers’ inner and outer surfaces for more application such as cellular therapies and drug- and gene-delivery strategies.

## Acknowledgments

This work was partially supported by the Office of Naval Research Grant N000141712620 and National Science Foundation Award 2014346.

## Data Availability Statement

The data supporting this study’s findings are available from the corresponding author upon reasonable request.

